# Shedding light on biogas: a transparent reactor triggers the development of a biofilm dominated by *Rhodopseudomonas faecalis* that holds potential for improved biogas production

**DOI:** 10.1101/521435

**Authors:** Christian Abendroth, Adriel Latorre- Pérez, Manuel Porcar, Claudia Simeonov, Olaf Luschnig, Cristina Vilanova, Javier Pascual

**Affiliations:** Robert Boyle Institut e.V., Jena, Germany.; Technische Universität Dresden, Chair of Waste Management, Pratzschwitzer Str. 15, Pirna, Germany.; Darwin Bioprospecting Excellence, S.L., Paterna, Valencia, Spain.; Institute for Integrative Systems Biology (I2SysBio), Paterna, Valencia, Spain.; Bio H2 Umwelt GmbH, Jena, Germany.

**Keywords:** *Anaerobic digestion*, *Rhodopseudomonas faecalis*, optimized biogas, shotgun metagenomic sequencing, waste water treatment, phototrophism, 16S rRNA gene amplicon Nanopore sequencing

## Abstract

Conventional anaerobic digesters intended for the production of biogas usually operate in complete darkness. Therefore, little is known about the effect of light on microbial communities operating in anaerobic digesters. In the present work, we have studied through 16S rRNA gene amplicon Nanopore sequencing and shotgun metagenomic sequencing the taxonomic and functional structure of the microbial community forming a biofilm on the inner wall of a lab-scale transparent anaerobic biodigester illuminated with natural sunlight. The biofilm was composed of microorganisms involved in the four metabolic processes needed for biogas production. The biofilm proved surprisingly rich in *Rhodopseudomonas faecalis*, a versatile bacterium able to carry out a photoautotroph metabolism when grown under anaerobic conditions. Our results suggest that this bacterium, able to fix carbon dioxide, could be considered for its use in transparent biogas fermenters in order to contribute to the production of optimized biogas with a higher CH_4_:CO_2_ ratio than the biogas produced in regular, opaque digesters. To the best of our knowledge, this is the first study supporting illuminated bioreactors as a new bioprocess for the obtention of biogas enriched in methane.

## 1. Introduction

Anaerobic digestion (AD) of organic matter is a robust technology for biogas synthesis from different types of waste (Borjesson and Mattiasson, 2008), and numerous studies have been conducted to optimise the synthesis of biogas and evaluate potential substrates (Jiang et al., 2018). Anaerobic digesters can be fed with a wide range of substrates such as grass biomass (Abendroth et al., 2017), sewage sludge from water treatment (Hardegen et al., 2018), microalgal biomass (Doloman et al., 2017), and food waste (Yang et al., 2016), among others. The main goal of AD is the production of biogas, a renewable energy that can be used for heating, electricity, and many other operations that use combustion engines (Lindkvist et al., 2017). Biogas is a mixture of methane (CH_4_; 55-70% of the total volume), carbon dioxide (CO_2_; 30-40%) and traces of other gases such as hydrogen sulfide (H_2_S) (Richards et al., 1994). Whereas methane is a flammable gas over a relatively large range of concentrations in air at standard pressure (5.4–17%), carbon dioxide is an inert gas. Therefore, increasing the CH_4_:CO_2_ ratio is one of the keystones for the production of high-quality biogas. The CH_4_:CO_2_ ratio can vary depending on the design of the biodigester, the substrate composition, as other factors such as temperature, pH, and substrate concentration (Hafner and Rennuit; 2017). An appropriate design of the anaerobic digestor is thus central for the production of optimized biogas.

The microbial communities operating in the digester are the final key players responsible for the quality of the produced biogas. The role of different microorganisms in the four metabolic steps carried out during the AD of organic matter (hydrolysis, fermentation, acetogenesis, and methanogenesis) has been widely studied (Zinder, 1984). A diverse number of *Bacteria* are known to be involved in the hydrolysis and further fermentation of complex polymers, whereas the oxidation of intermediate fermentation products to acetate is performed by either hydrogen- or formate-producing acetogens (Stams & Plugge, 2009). Lastly, methane synthesis is mainly derived from acetate and H_2_/CO_2_ by acetoclastic and hydrogenoclastic methanogenic *Archaea*. Therefore, an improved understanding of the microbial communities and their metabolic roles during the four stages of AD may also help to optimize biogas production in terms of quantity (yield) and quality (CH_4_:CO_2_ ratio of the produced gas). Over the past few years, next-generation sequencing techniques, such as 16S rRNA gene amplicon sequencing and shotgun metagenomic sequencing, have been applied to study the structure and composition of microbial communities in different types of anaerobic digesters (Hanreich et al., 2013; Sundberg et al., 2013; Maus et al., 2016; Abendroth et al., 2015; Abendroth et al., 2017a; Hardegen et al., 2018). These studies have shown that each type of bioreactor harbors a specific microbial community (Sundberg et al., 2013; Abendroth et al., 2015). Each particular community is determined by parameters such as the type of feedstock (Sundberg et al., 2013), temperature (Banach et al., 2018), retention time (Gaby et al., 2017), salt content (Wang et al., 2017; De Vrieze et al., 2017), viscosity (Hardegen et al., 2018), pH (Zhou et al., 2016), or the loading rate (Ciotola et al., 2014). Although the influence of many physicochemical parameters on microbial communities has been studied in anaerobic digesters, there is, to the best of our knowledge, no previous reports characterising the influence of light in the process (Sawayama, 2000; Tada et al., 2006; Wei et al., 2016; Reverso, 2017), mainly because the obvious fact that conventional AD systems operate in complete darkness. Interestingly, a previous study reported an increment of the relative concentration of methane when an anaerobic digester was operated under the influence of light (Tada et al., 2006). However, a holistic study of the effect of light on the entire microbiome of anaerobic digesters had yet to be addressed.

The aim of the present work was to analyse the effect of natural sunlight on the microbial community of a lab-scale anaerobic co-digester, in order to explore the potential of this strategy to produce high quality biogas. In order to reach that goal, we used full-length 16S rRNA gene amplicon Nanopore sequencing, shotgun metagenomic sequencing and a complete bioinformatics analysis to unveil the structure and composition of the microbial community grown as a red-coloured biofilm over the transparent wall of a specifically designed transparent leach-bed bioreactor.

## 2. Material and Methods

### 2.1 Substrate and seed sludge

Untreated grass biomass (*Graminidae*) from a pasture in Jena (Germany) was used as feedstock. Collected grass biomass was characterised by a total solids content (TS) of 30.4%, with 84.2% of the TS being volatile solids (VS). TS and VS were determined as described in Abendroth et al. (2017). One gram of fresh biomass showed a chemical oxygen demand (COD) of 260 mg O_2_/L. As seed sludge, sewage sludge from an anaerobic digester of the water treatment plant in Jena (Germany) was used.

### 2.2 Digestion conditions

The experiment was carried out in an open hall during summer of 2017. A transparent two-stage leach system of plexiglas^®^ was built, and used to perform acidification of grass biomass in a leach-bed configuration and methanisation using an anaerobic filter as described in Abendroth et al. (2017). The anaerobic digester was placed in a location where it received indirect natural sunlight (ca. 10 hours daylight). The anaerobic digester was placed in a location where it received indirect natural sunlight (ca. 10 hours daylight). The acidification stage and the methane stage had a working volume of 20 L each and both stages were treated at mesophilic temperature (37 °C). Grass biomass was retained in a strainer during acidification. The methane stage was filled with 1.58 kg of bed packing (Hel-X, Christian Stöhr, Germany). Both stages were filled with sewage sludge at the beginning of the experiment: 8 L for the acidification stage and 11.75 L for the methane stage. For every batch cycle of acidification, 96.2 gL^−1^ of fresh grass biomass were used. During acidification, the pH was kept at 6.0 using a pH-regulation system (BL 7916, Hanna Instruments, Germany). After each cycle, the collected liquor was stored at 4 °C. Each methane stage received daily approximately 100 gCOD (8.5 gCOD L−1). Gas production was quantified with a customized gas counter and collected in gasbags (Tecobag, Tesseraux, Germany) for further analysis.

### 2.3 Sample collection and metagenomic DNA isolation

The anaerobic digester was operated for three weeks prior to sampling. Three independent biofilm samples (ca. 250 μl each) were collected by scratching the inner bioreactor wall and further mixed in an Eppendorf tube. In order to reduce the amount of inhibiting substances, the biofilm sample was centrifuged (10 min at 20,000 *g*) and then washed several times with sterile phosphate-buffered saline (PBS) pH 7.2 until a clear supernatant was observed. Subsequently, metagenomic DNA was performed using the Power Soil DNA Isolation kit (MO BIO Laboratories) following the manufacturer’s instructions. DNA concentration, quality and integrity were assessed with a Nanodrop-1000 spectrophotometer (Thermo Scientific, Wilmington, DE) and on a 1.5% (w/v) agarose gel, respectively.

### 2.4 Full-length 16S rRNA gene amplicon Nanopore sequencing

The bacterial full-length 16S rRNA gene was amplified via PCR using the primer pair S-D-Bact-0008-a-S-16 (5’-AGRGTTYGATYMTGGCTCAG-3’) and S-D-Bact-1492-a-A-16 (5’-TACCTTGTTAYGACTT-3’) (Klindworth et al., 2013). The following reagents and concentration were used for the first PCR reaction: 200 μM dNTPs, 200 nM of each primer, 1 U of VWR Taq DNA Polymerase (VWR^®^, WR International bvba/sprl, Belbium), 1 X key Buffer supplemented with MgCl_2_ (1.5 mM), and 10 ng of DNA template (final volume: 50 μL). PCR started with an initial denaturation step at 94°C for 1 min, followed by 35 cycles of amplification (denaturing, 1 min at 95°C; annealing, 1 min at 49°C; extension, 2 min at 72°C), and a final extension at 72°C for 10 min. A negative control without DNA template was included. Agencourt AMPure XP beads (Beckman Coulter) at 0.5X concentration were used to remove primer-dimers and non-specific amplicons. DNA concentration was measured using the Qubit™ dsDNA HS Assay kit (Qubit^®^ 2.0 Fluorometer, Thermofisher, Waltham, USA). Then, the Ligation Sequencing Kit 1D (SQK-LSK108) was used to prepare the amplicon library to load into the MinION™. The flow cell (R9.4, FLO-MIN106) was primed and then loaded as indicated in the ONT™ protocols.

Reads were basecalled in real time using the MinKNOW™ software (version 1.13.1, standard sequencing protocol), and sequencing statistics were followed in real time using the EPI2ME debarcoding workflow. Porechop (https://github.com/rrwick/Porechop) was applied for removing the adaptors. By default, MinKNOW™ software removed reads with quality values lower than 7 in the PHRED score. The resulting sequences were analysed using the QIIME (Kuczynski *et al.*, 2011). Briefly, reads were taxonomically classified through BLAST searches against the latest version of the GreenGenes database (13.8; DeSantis *et al.*, 2006). Rarefaction curves of the full-length 16S rRNA reads, including and excluding singletons, were obtained using iNEXT (v. 2.0.17) R package.

### 2.5 Shotgun metagenomic sequencing

The biofilm sample was also subjected to shotgun metagenomic sequencing. Briefly, the Nextera XT Prep Kit protocol was followed for library preparation. Then, Illumina MiSeq^®^ platform was used for sequencing. The parameters were adjusted to obtain pair-end sequences of 150 bp and a sequencing depth of 10 million of reads. Adapters were trimmed, and a quality filtering was applied with BBDuk (included in BBTools package; Bushnell B., https://sourceforge.net/projects/bbtools/ updated January 2, 2018). Reads shorter than 50 bp and/or with a mean quality lower than Q20 (in PHRED scale) were discarded. The quality parameters of the sequences were checked with FastQC (http://www.bioinformatics.babraham.ac.uk/projects/fastqc, version 0.11.7).

The quality-checked reads were taxonomically classified via Centrifuge (v. 1.0.3; Kim *et al.*, 2016) against a compressed database containing reference sequences from *Bacteria* and *Archaea* (updated April 15, 2018; available at https://ccb.jhu.edu/software/centrifuge/). The shotgun sequences were assembled into contigs and scaffolds with the metaSPAdes pipeline included in SPAdes (v. 3.12.0; Nurk *et al.*, 2017). The statistics and attributes of the assembly were explored with QUAST (v. 4.6.3). Selected scaffolds were grouped into bins with MaxBin2 (v. 2.2.4; Wu *et al.*, 2015) in order to reconstruct metagenome-assembled genomes (MAGs). CheckM (v1.0.11; Parks *et al.*, 2015) was used for assessing the quality and completeness of each MAG. Only high-quality MAGs, with completeness values greater than 50% and contamination values lower than 10%, were considered for further analyses.

The taxonomic affiliation of each MAG was assessed with the *Similar Genome Finder Service* tool of the PATRIC (Wattan et al., 2017). This tool matched each MAG against a set of representative and reference genomes available in PATRIC (Wattan et al., 2017) by using Mash/MinHash distances (Ondov et al., 2016). Subsequently, a phylogenomic tree was inferred for each MAG in order to know their specific evolutionary history. The UBCG v. 3.0 pipeline (Up-to-date bacterial core gene set; Na *et al.*, 2018) was used to construct maximum likelihood trees based on a multiple alignment of a set of 92 to 90 universal and single copy gene sequences (Table S1). Despite the fact that the UBCG pipeline is optimized for bacterial genomes, it was also used for archaeal MAGs but using 23 universal and single copy genes. In order to investigate if each MAG belonged to a known species, pairwise average nucleotide identity values (ANIb) (Goris et al., 2007) were calculated between each MAG and its closest type strain, by using the JSpeciesWS online tool (Richter et al., 2015). Additionally, digital DNA-DNA hybridization (DDH) pairwise values were also obtained using the Genome-to-Genome Distance Calculator 2.1 (GGDC) tool (Meier-Kolthoff et al., 2013). As recommended for incompletely sequenced genomes, Formula 2 was used for calculating the digital DDH values (Meier-Kolthoff et al., 2013).

### 2.6 Functional analysis of the microbial community

The assembled metagenomic sequences, as well as the high-quality MAGs, were annotated using the RAST toolkit (RASTtk) (Brettinet al., 2015) implemented in the Genome Annotation Service in PATRIC (Wattam et al., 2017). The carbohydrate-active enzymes (CAZyme) of each MAG were determined by identifying genes containing CAZymes domains using the dbCAN2 meta server (Zhang et al., 2018). CAZyme domains were predicted by HMMER (E-Value < 1e-15, coverage > 0.35). The metabolic pathways reconstruction for each MAG was done comparing the protein-coding genes against the Kyoto Encyclopedia of Genes and Genomes (KEGG) (Kanehisa et al., 2017) and MetaCyc metabolic pathways (Caspi et al., 2018).

## 3. Results and Discussion

### 3.1. Taxonomic diversity of the microbial community

After three weeks operating the anaerobic digester, a bright, red-pigmented microbial biofilm appeared on the inner wall of the bioreactor (Fig. 1). Since the microbiome of the main content of the biodigesters has already been extensively studied (Hanreich et al., 2013; Sundberg et al., 2013; Maus et al., 2016; Abendroth et al., 2015; Abendroth et al., 2017; Hardegen et al., 2018), we focused on the characterisation of the microbiome of the red-pigmented biofilm developed on the transparent wall of the bioreactor.

**Fig. 1.**
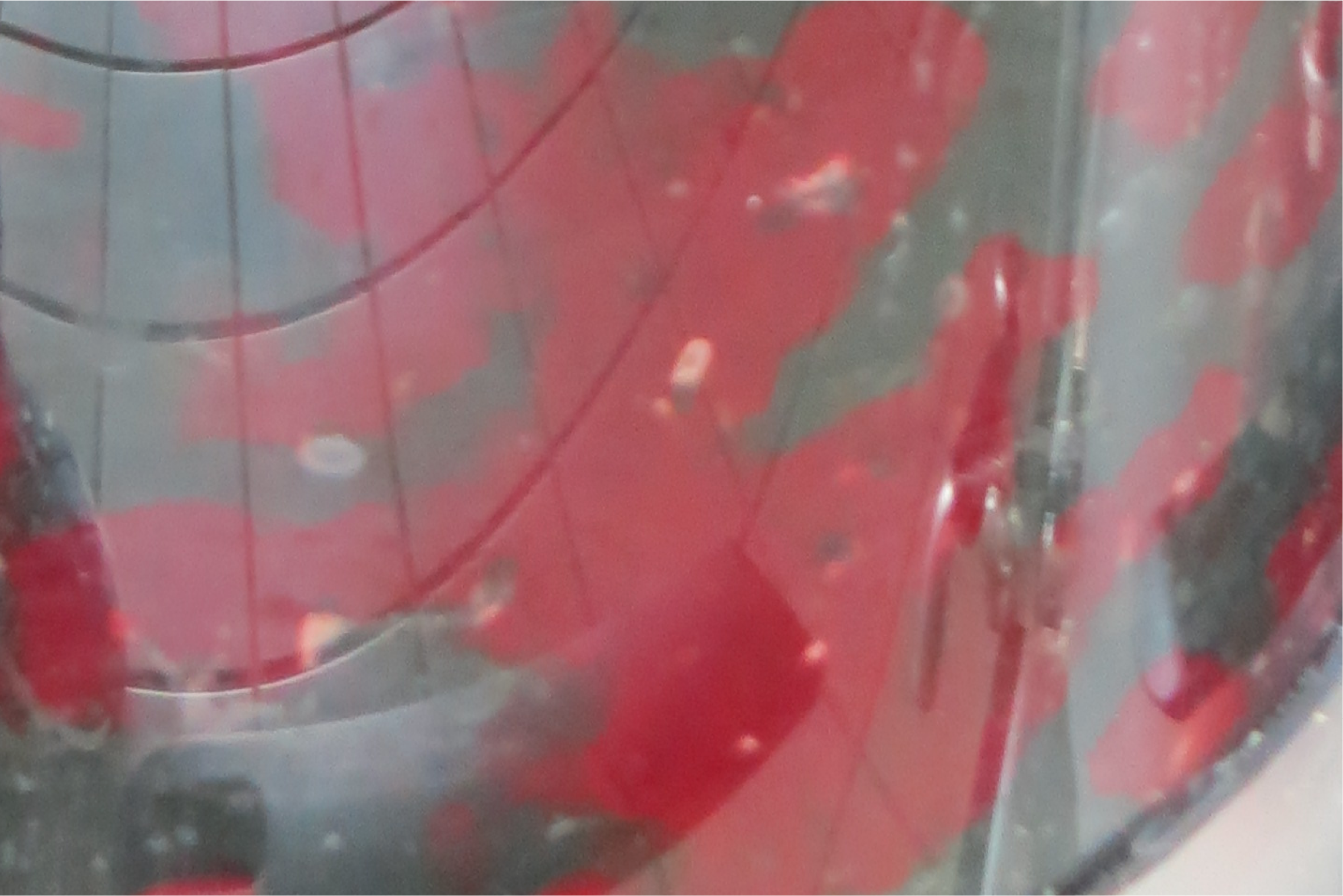
Photograph of the transparent wall of the anaerobic digester used to analyse the effect of natural sunlight on the microbial community. After three weeks operating de anaerobic digester, a red-pigmented microbial biofilm grew on the inner wall of the reactor forming irregular patches.

The taxonomic profile of the microbiome of the biofilm was studied via 16S rRNA amplicon Nanopore sequencing as well as by sequencing the whole 16S rRNA gene without the need for *in silico* assembly (Kerkhof et al., 2017). After a quality filtering of the raw reads, a total of 11,163 16S rRNA sequences were retrieved and taxonomically classified. The median sequence length was 1,445 nt and the mean read quality was 9.8. Similarly to Ma et al. (2017), any attempt of clustering the reads into Operational Taxonomic Units (OTUs) using the closed-reference OTU picking method available in Qiime failed (i.e. each sequence was classified as an independent OTU). The number of reads obtained was enough to analyse the vast majority of the microbial species (Fig. S1A), thus enabling a comprehensive characterization of the microbial community. The saturation of the species richness was more evident when the singletons were excluded from the dataset (Fig. S1B).

The bacterial community was dominated by members of the phyla *Firmicutes, Bacteroidetes* and *Proteobacteria*, followed by *Chloroflexi, Spirochaetes* and the candidate phylum WS6 (Fig. 2A). Additionally, members of 43 phyla or candidate divisions were also identified (Table S1). This profile was similar to that found by other authors in dark AD (Hanreich et al., 2013; Sundberg et al., 2013; Maus et al., 2016; Abendroth et al., 2015; Abendroth et al., 2017). At the family level, *Gracilibacteraceae, Lachnospiraceae* and *Tissierellaceae* were the dominant *Firmicutes*, while *Porphyromonadaceae* and *Bradyrhizobiaceae* were the most abundant *Bacteroidetes* and *Alphaproteobacteria*, respectively (Fig. 2A; Table S1). Additionally, 36 reads were classified as *Archaea.* However, *s*ince the primer pair used to amplify the 16S rRNA was optimized for *Bacteria* (Klindworth et al., 2013), all archaeal reads were excluded from the analysis.

**Fig. 2.**
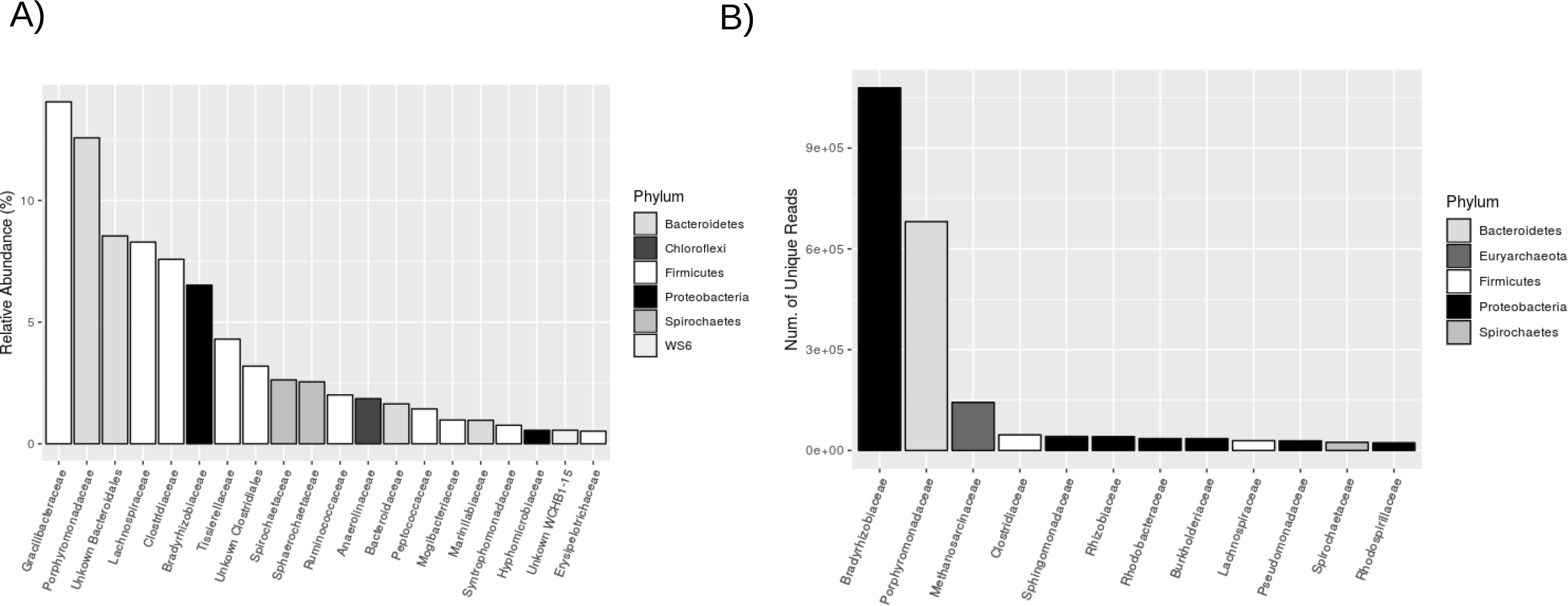
(A) Taxonomic classification of the full-length 16S rRNA reads sequenced with the Nanopore technology. Reads were blasted against the GreenGenes database (v 13.8). Only those families with a relative abundance higher than 0.5% are shown. (B) Taxonomic classification of the shotgun metagenomic Illumina reads. Reads were mapped against a database containing reference sequences from *Bacteria* and *Archaea* (updated April 15, 2018; available at https://ccb.jhu.edu/software/centrifuge/). Only the top 12 most abundant families are shown.

Although the full-length sequence of the 16S rRNA gene was sequenced, a high number of phylotypes could only be classified up to the family level (Table S1), suggesting a high taxonomic novelty of the taxa. The high diversity of phylotypes recovered from the biofilm sample (723 phylotypes, Table S1), might be a consequence of the high error rate of the Nanopore sequencing technology. In fact, 46.6% of microbial phylotypes were singletons and nine phyla were represented by a single read (Table S1). Nevertheless, a study has recently demonstrated that the MinION technology has the ability to provide rRNA operon sequence data of sufficient quality for characterizing the microbiota of complex environmental samples and provides results that are reproducible, quantitative and consistent (Kerkhof et al., 2017). Another explanation why a high number of phylotypes were identified in the biofilm is that the seed sludge and the feedstock used was carrying a highly diverse microbial load (Kirkegaard et al., 2017), albeit those communities might not be metabolically active in the sampled biomass. An indication of the presence of inactive microorganisms in the community is the occurrence of obligate aerobic bacteria, such as *Arthrobacter* or *Devosia* (Table S1). Further studies based on metagenomic RNA would be necessary to distinguish between the phylotypes that are keyplayers in the biofilm and those that are merely transported by the influent as inactive microorganisms.

In order to complement the taxonomic information of the microbial community of the red-coloured biofilm, shotgun metagenomic sequencing was also performed. A total of 8,903,087 high-quality pair-end reads with a median size of 150 nt were sequenced. The taxonomic classification of all the metagenomic reads is shown in Fig. 2B. Only 38.2 % of metagenomic reads mapped against the genomic database, corroborating the taxonomic novelty of the microorganisms that form the biofilm. Most of the reads mapped against genomes of *Bradyrhizobiaceae*, specifically *Rhodopseudomonas palutris* (26.6% of total reads), followed by the *Porphyromonadaceae* species *Fermentimonas caenicola* (22.9%) and the archaeal species *Methanosarcina mazei* (4.7%). Interestingly, and in sharp contrast with what has been described for regular (dark) anaerobic digesters (Hanreich et al., 2013; Sundberg et al., 2013; Maus et al., 2016; Abendroth et al., 2015; Abendroth et al., 2017; Hardegen et al., 2018), the illumination of the bioreactor triggered an enrichment of *Rhodopseudomonas* (Fig. 2B). Similarly, other authors reported enrichment of *Rhodopseudomonas faecalis* in an illuminated anaerobic digester fed with swine sewage wastewater (Wei et al., 2016). In contrast to 16S rRNA sequencing results, the family *Gracilibacteraceae* was not abundant in the shotgun metagenomic data (Fig. 2). This type of taxonomic discrepancies between both sequencing approaches has previously been discussed by other authors (Tessler et al., 2017).

### 3.2. Functional profile of the community and definition of microbial keyplayers

A total of 6183 contigs composed by 38,667,755 nt were assembled from the shotgun metagenomic sequencing. 40,136 coding sequences (CDS) were identified after the functional annotation of contigs with RASTtk. 54.9% of CDS were identified as proteins with functional assignments, while the remaining CDS were annotated as hypothetical proteins. Furthermore, 556 tRNA, 53 rRNA, 583 CRISPR-repeats, 556 CRISPR-spacers and 27 CRISPR-array sequences were also reported. 37% of the CDS were assigned to functional subsystems (Fig. 3; Table S3). The great majority of the CDS (36.8%) were involved in cellular metabolism, including genes engaged in the turnover of nutrients (Table S3). Genes related to protein processing accounted for 17.3%, while those for energy and DNA processing accounted for 12.4% and 6.5%, respectively (Fig. 3).

**Fig. 3.**
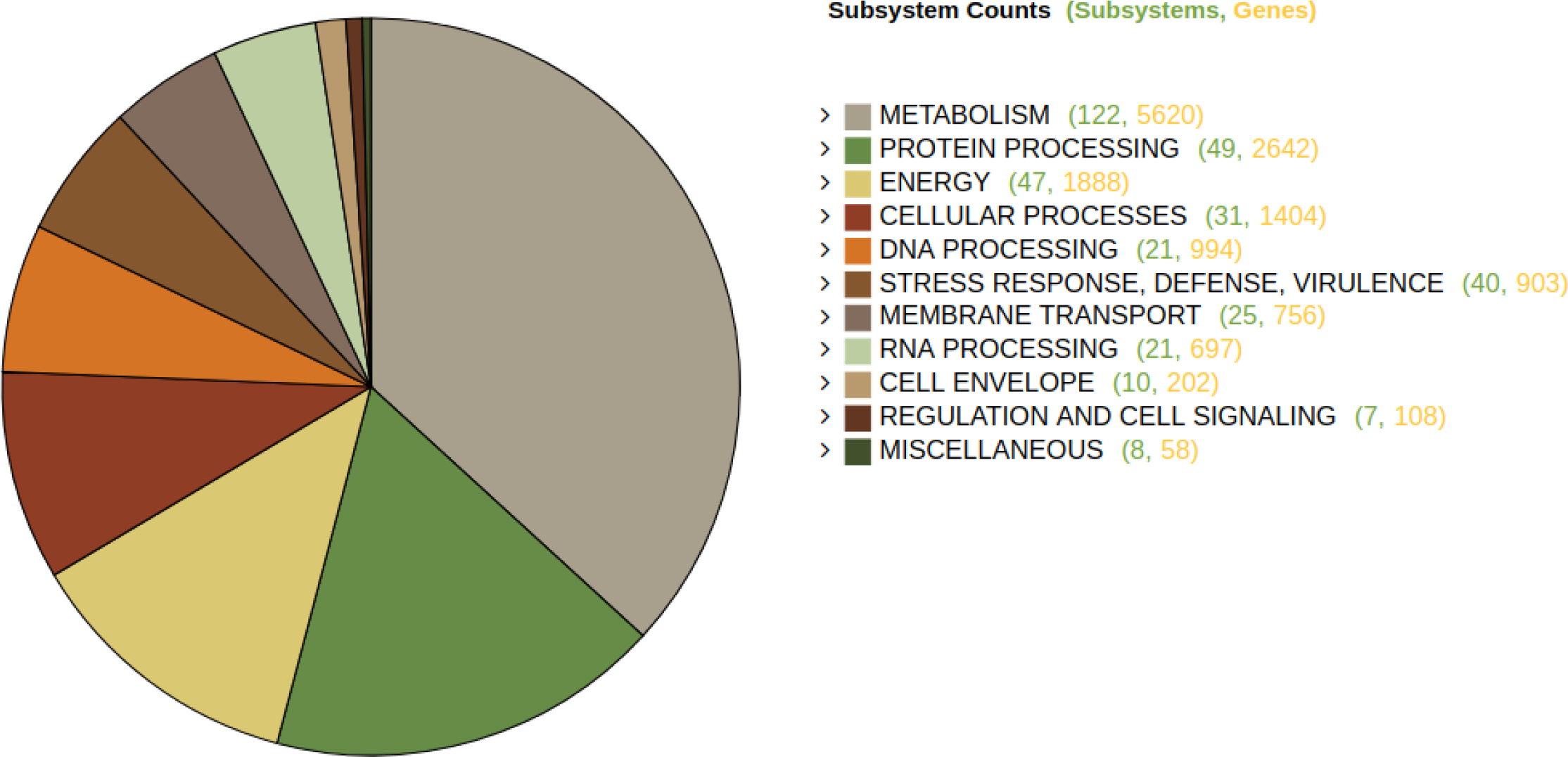
Classification of annotated coding sequences (CDS) in functional subsystems.

To date, many microbial ecology studies in anaerobic digesters have been based on 16S rRNA OTUs. However, due to the great metabolic diversity of certain taxa as well as the impossibility to classify some OTUs at lower taxonomic levels, it is difficult to predict the accurate functional roles that each microorganism plays during the AD (Kirkegaard et al., 2017). Therefore, in order to shed light on the key microorganisms and their metabolic functions in anaerobic digesters when operated under the influence of natural sunlight, assembled contigs were binned as MAGs. Eight out of the 24 MAGs obtained passed the filter of contamination and completeness (Table 1). The number of scaffolds of the eight high-quality MAGs ranged from 141 to 985 and the estimated genome size, from 1.2 Mb to 4.0 Mb (Table 1). The G+C content of four MAGs was equal to or less than 37.0% and none of them showed a value greater than 64.2% (Table 1). MAG 9 harboured chimeric ribosomal RNA operons and hence their 16S rRNA gene sequences could not be used for taxonomic purposes, while other MAGs like MAG 16 did not include any ribosomal RNA operon. The MAG 1 was identified as a member of the species *Rhodopseudomonas faecalis* (Table 2; Fig. S1). MAG 2 was identified as a strain of the *Fermentimonas caenicola* species, while MAG 16 was identified as *Methanosarcina mazei*. The ANI and digital DDH values between MAGs 1, 2 and 16 and the type strains of phylogenetically close species were higher than the threshold established to circumscribe prokaryotic species, namely 95% for ANI values (Richter and Rosselló-Móra, 2009) and 70% for digital DDH (Meier-Kolthoff et al., 2013). Therefore, both genome-related indexes (Chun and Rainey, 2014) confirmed the adscription of MAGs 1, 2 and 16 to previously known species. Based on the number of reads obtained from the shotgun metagenomic sequencing, MAG 1 (*Rhodopseudomonas faecalis)* and MAG 2 (*Fermentimonas caenicola*) were the most abundant bacteria in the red-pigmented biofilm; while MAG 16 (*Methanosarcina mazei*) was the only archaeon identified in the community (Fig 2). Contrarily, the other five MAGs could not be identified at the species level, but at the genus level or even a higher taxonomic rank (Table 2). MAG 4 was classified as a new taxon of the family *Anaerolinaceae* (phylum *Chloroflexi*), being closely related to the strain *Anaerolinea* sp. CP2_2F, a bacterium recently isolated from a methanogenic waste water treatment system (Matsuura et al., 2015). Two MAGs were classified as members of the *Spirochaetaceae* family. Specifically, MAG 5 was closely related to *Sphaerochaeta globosa* Buddy^T^, a strain isolated from an anoxic river sediment (Ritalahti et al., 2012), while MAG 11 was identified as a new species of the genus *Treponema*. MAG 10 was classified as a novel lineage of the family *Gottschalkiaceae*, being phylogenetically related to the *Gottschalkia purinilytica* DSM 1384^T^ (Poehlein et al., 2017). Finally, MAG 9 was classified as a hitherto unknown taxon of the family *Erysipelotrichaceae* (phylum *Firmicutes*).

**Table 1:** Summary statistics of the reconstructed metagenome-assembled genomes (MAGs) used in this study. The completeness and contamination of each MAG were estimated with CheckM (Parks *et al.*, 2015) and the coverage with MaxBin2 (Wu et al., 2015).

| MAGs | Contigs | Genome length (Mb) | Coverage | Completeness (%) | Contamination (%) | G+C (%) | CDS | Proteins with functional assignments | Hypothetical proteins |
| --- | --- | --- | --- | --- | --- | --- | --- | --- | --- |
| 1 | 119 | 4.0 | 76.5 | 98.4 | 0.17 | 64.2 | 3797 | 2725 | 1072 |
| 2 | 141 | 2.7 | 31.8 | 100 | 0.55 | 37.0 | 2348 | 1529 | 819 |
| 4 | 319 | 3.2 | 17.8 | 91.2 | 0.67 | 37.0 | 2946 | 1821 | 1125 |
| 5 | 136 | 2.2 | 10.9 | 92.0 | 3.45 | 53.7 | 2270 | 1266 | 1004 |
| 9 | 213 | 1.2 | 6.8 | 96.6 | 4.96 | 36.3 | 1191 | 585 | 606 |
| 10 | 352 | 1.6 | 6.7 | 53.5 | 5.2 | 31.6 | 1792 | 1027 | 765 |
| 11 | 346 | 2.7 | 5.7 | 73.4 | 3.26 | 53.2 | 2770 | 1297 | 1473 |
| 16 | 985 | 2.8 | 4.2 | 81.0 | 4.63 | 42.9 | 3459 | 2094 | 1365 |

**Table 2.**
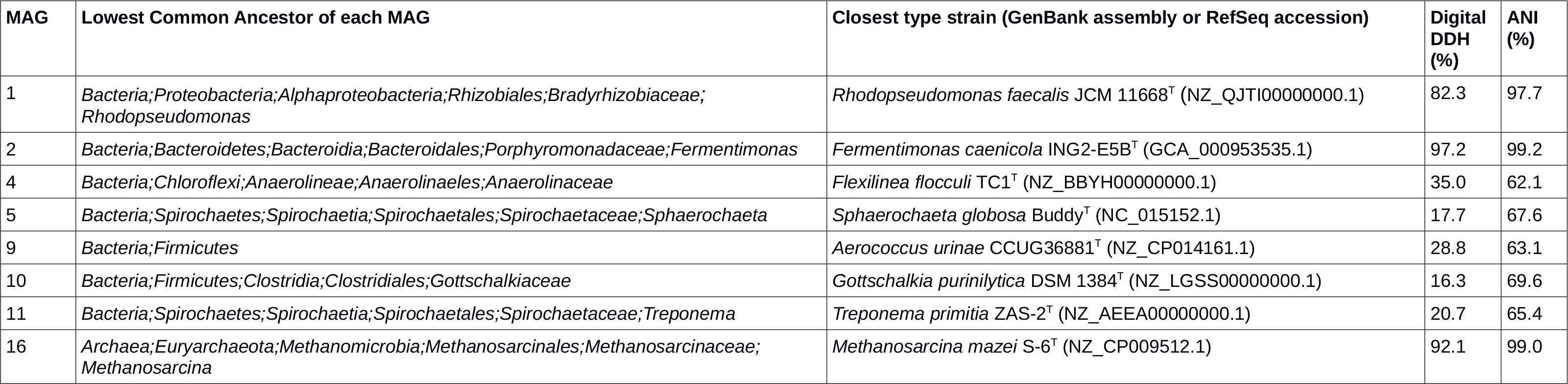
Lowest common ancestor and closest type strain of each metagenome-assembled genome (MAG). The average nucleotide identity (ANI) and digital DNA-DNA hybridization (DDH) values with regard to its closest type strain are shown for each MAG.

### 3.3. Affiliation of functional CDS to the four stages of anaerobic digestion

#### 3.3.1 Hydrolysis of complex polymers

A total of 108 different glycoside hydrolases families (Carbohydrate-Active enZYmes Database; Lombard et al., 2014) were found in the eight high-quality MAGs (Fig. 4; Table S4). The microorganism with a greater repertoire of glycoside hydrolases was MAG 4 (*Anaerolinaceae* sp.), which contains 104 glycoside hydrolases of over 49 families. MAG 2 (*Fermentimonas caenicola*) codified for 82 glycoside hydrolases of 41 different families; and MAG 11 (*Treponema* sp.) encoded 54 glycoside hydrolases belonging to 36 families. Furthermore, MAGs 4, 2 and 3 were also the major producers of carbohydrate esterases (Fig. 4; Table S4). Representatives of glycosyl transferase families GT2_Glycos_transf_2 and GT4 were found in the eight MAGs. Moreover, MAG 4 contained a high number of GT2_Glycos_transf_2-coding genes, specifically 18 protein-coding genes (Fig. 4; Table S4). Only a single polysaccharide lyase, encoded by MAG 5, was detected in the whole metagenome.

**Fig. 4.**
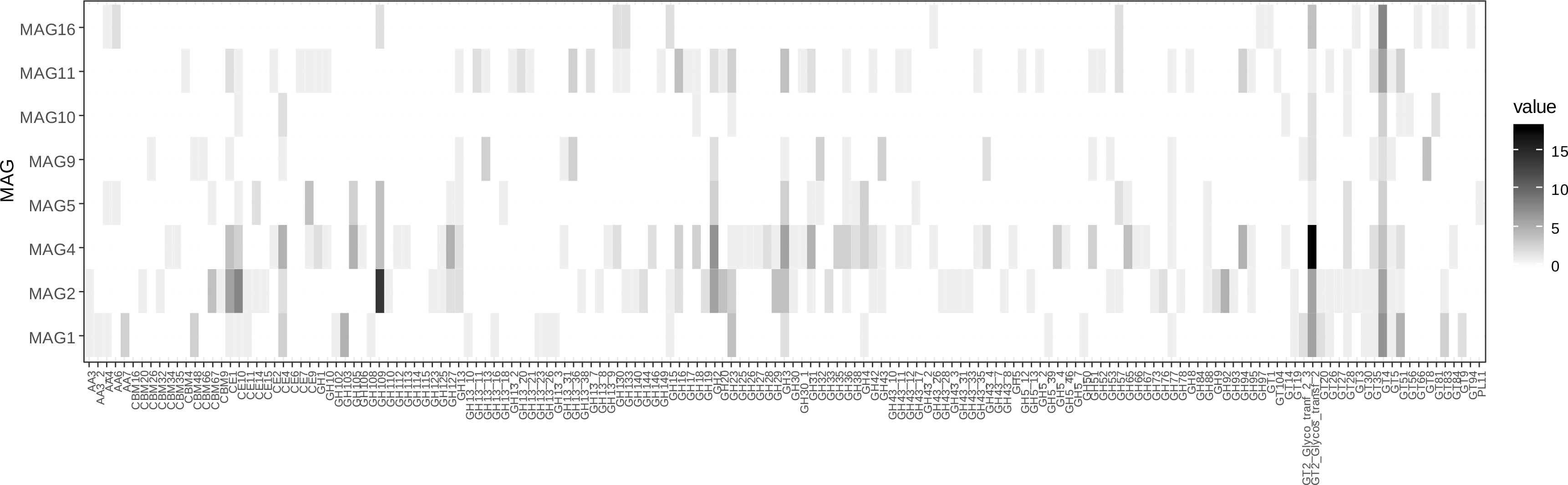
Heatmap of Carbohydrate-Active Enzymes (CAZymes) families found in each of the eight metagenome-assembled genomes. Both CAZY families and MAGs are listed alphabetically. The color intensity corresponds to the number of protein-coding genes identified in each family. CAZyme family codes: GT, glycosyltransferases; GH, glycoside hydrolases; CE, carbohydrate esterases; PL, polysaccharide lyases; CBM, carbohydrate binding modules; AA, axillary activities (oxidative enzymes).

The nature of the substrate determines the type of bacteria involved in the hydrolysis step. Preeti Rao et al. (1993), observed that digesters fed with cow manure supported more amylolitic microorganisms, whereas digesters fed with poultry waste showed higher proteolytic populations. Since our anaerobic digester was fuelled with untreated grass biomass, a dominance of cellulolytic, hemicellulolytic and lignolytic bacteria such as *Chloroflexi*, *Bacteroides*, *Spirochaetes* and *Clostridium* was expected (Pinnell et al., 2014).

#### 3.3.2 Acidogenesis

Except for MAG 9 (*Erysipelotrichaceae)* and MAG 16 (*Methanosarcina mazei)*, all the other MAGs showed a potential fermentative metabolism based on a functional analysis of their genomes (Table 3). MAG 1 (*Rhodopseudomonas faecalis)*, 10 (*Gottschalkiaceae* sp.) and 11 (*Treponema* sp.) were able to carry out the fermentation of pyruvate to lactate with lactate dehydrogenase enzymes (Table 3; Table S5). MAGs 2 and 5 were able to conduct the transformation of pyruvate to (R)-2-acetoin thorough acetolactate synthase (EC 2.2.1.6) and acetoin dehydrogenase E1 (EC 2.3.1.190) enzymes (Table S5). The presence of the acetolactate synthase enzyme (EC 2.2.1.6) was also observed in MAGs 1, 4 and 11. MAG 2 was able to generate (R, R)-2,3-butanediol from (R)-2-acetoin with the 2,3-butanediol dehydrogenase enzyme (EC 1.1.1.4). MAG 11 encoded two acetaldehyde dehydrogenases (EC 1.2.1.10), while MAG 4 encoded alcohol dehydrogenase (EC 1.1.1.1) and pyruvate formate-lyase (EC 2.3.1.54) enzymes, allowing the transformation of acetaldehyde to ethanol and pyruvate to formate, respectively.

**Table 3:** Summary of metabolic pathways identified in each metagenome-assembled genome (MAG). ND, not detected.

| Pathway | MAG 1 | MAG 2 | MAG 4 | MAG 5 | MAG 9 | MAG 10 | MAG 11 | MAG 16 |
| --- | --- | --- | --- | --- | --- | --- | --- | --- |
| Fermentation | Active | Active | ND | Active | ND | Active | ND | ND |
| Homoacetogenesis<sup>a</sup> | ND | ND | ND | ND | ND | ND | Active | Active |
| Acetogenesis | ND | Active | Active | ND | ND | ND | ND | ND |
| Methanogenesis from acetate | ND | ND | ND | ND | ND | ND | ND | Active |
| Methanogenesis from CO<sub>2</sub> and H<sub>2</sub> | ND | ND | ND | ND | ND | ND | ND | ND |
| Methanogenesis from methanol | ND | ND | ND | ND | ND | ND | ND | ND |
| Calvin-Benson cycle<sup>b</sup> | Active | ND | ND | ND | ND | ND | ND | ND<sup>b</sup> |
| Photosystem II-type photosynthetic reaction center | Active | ND | ND | ND | ND | ND | ND | ND |
| Nitrogen fixation | Active | ND | ND | ND | ND | ND | ND | ND |

<sup>a</sup> Reductive acetyl-CoA or Wood-Ljungdahl pathway
<sup>b</sup> Type III ribulose-1,5-bisphosphate carboxylase-oxygenase (RuBisCO) is present.

#### 3.3.3 Acetogenesis

In our anaerobic digester operating under the influence of natural sunlight, *Methanosarcina* (MAG 16) was the only microorganism harboring the genes corresponding to the Wood-Ljungdahl pathway (Table 3; Table S6). The Wood–Ljungdahl pathway coupled to the methanogenesis is one of the most ancient metabolisms for energy generation and carbon fixation in *Archaea* (Borrel et al., 2016). The alternative acetogenesis pathway (acetogenesis by dehydrogenation or syntrophic acetogenesis) is based on the anaerobic oxidation of long and short chain (volatile) fatty acids. MAGs 2 (*Fermentimonas caenicola*) and 4 (*Anaerolinaceae* sp.) contained some key genes involved in the conversion of propionate to acetate (Table 3; Table S5). Additionally, MAG 4 and MAG 11 (*Treponema* sp.) contained the enzymes necessary to perform the conversion of acetyl-CoA to acetate, namely acetate kinase (EC 2.7.2.1) and phosphate acetyltransferase (EC 2.3.1.8).

#### 3.3.4 Methanogenesis

The amount of methane produced in the anaerobic digester illuminated with natural sunlight ranged was approximately 400 ml/gCOD L^−1^. Interestingly, *Methanosarcina* (MAG 16) was the only methanogenic archaeon detected in the red-pigmented biofilm (Fig. 2B). MAG 16 harboured all the protein-coding genes in the acetoclastic pathway for methane production, except tetrahydromethanopterin S-methyltransferase (EC 2.1.1.86) and methyl-coenzyme M reductase (EC 2.4.8.1). However, both genes were also found in the whole metagenome and were identified as closely related to *Methanosarcina mazei* S-6 (NZ_CP009512.1). Despite that *Methanosarcina* has also been described as a hydrogenoclastic methanogen (De Vrieze et al., 2012), the key genes required for the utilization of H_2_ and CO_2_ for methane production were not found in MAG 16 (Table S6). Previous research found that approximately 70% of the methane produced in the digestion of domestic sludge comes from the transformation of acetate to methane by the acetoclastic methanogens (Jeris and McCarty, 1965). Usually, *Methanothrix* (formerly named *Methanosaeta)* and *Methanosarcina* co-exist in the anaerobic digesters and their relative abundances are driven by the acetate concentration (Conklin et al., 2006). *Methanosarcina* has greater rates of acetate utilization and growth, and greater half-saturation and yield coefficients compared to *Methanothrix* (Conklin et al., 2006). Therefore, a possible cause for the dominance of *Methanosarcina mazei* over *Methanothrix* species in the illuminated anaerobic digester may be the high concentration of acetate in the biomass. *Methanosarcina* is as very robust methanogen able to adapt to environmental changes (De Vrieze et al., 2012), as well as an efficient methane producer (Tada et al., 2006; De Vrieze et al., 2012). Indeed, a previous study assessing the effect of light on methane production during AD reported an enrichment of *Methanosarcina* spp. coupled with an increment in methane production (Tada et al., 2006). This suggests, in concordance with our results, that anaerobic digesters operated under light conditions may results in an enrichment of *Metanosarcina*. Direct interspecies electron transfer (DIET) is a process that takes place during AD and is of great importance, as it may significantly accelerate methanogenesis (Kato et al., 2012; Kato et al., 2018). It can be enhanced by adding electrically conductive particles. So far, *Methanothrix* and *Methanosarcina* are the only genera where DIET has been proved. However, since *Methanothrix* typically grows at acetic acid concentrations lower then 3000 mg L^−1^ (De Vrieze, 2014), *Methanosarcina* remains as the only known methanogen, which can apply DIET within high-performance digesters with high COD concentrations. Therefore, the illumination of anaerobic digesters might enhance the application of DIET due to its enrichment of *Methanosarcina*.

### 3.4. Functional novelty and implications

Unlike conventional anaerobic digesters operating in complete darkness, an enrichment of *Rhodopseudomonas faecalis* (MAG 1) took place when the anaerobic digester was illuminated with natural sunlight. *R. faecalis* is a common purple non-sulfur (PNS) bacterium able to carry out a wide range of metabolic pathways (Boone and Mah, 2015). MAG 1 has the entire repertoire of enzymes needed to synthesize a photosystem II-type photosynthetic reaction center (Table 3; Table S7). Photosynthetic reaction centers are complexes composed of several proteins, pigments and other co-factors that, together, execute the primary energy conversion reactions of photosynthesis. The pigments produced by *Rhodopseudomonas palustris*, closely related to *R. faecalis*, are both bacteriochlorophyll(BChl)-a and bacteriopheophytin(BPhe)-a (Mizoguchi et al., 2012). Therefore, the red to brownish-red colour of the biofilm developed in the bioreactor is very likely due to the massive presence of *Rhodopseudomonas faecalis* in the biofilm, although analysis with liquid chromatography-mass spectrometry would be necessary in order to confirm this hypothesis. MAG 1 is able to carry out the CO_2_ fixation via the Calvin-Benson cycle under anaerobic conditions (Table S7). In this process, CO_2_ and ribulose bisphosphate (5-carbon sugar) are transformed into 3-phosphoglycerate (Zheng et al., 2018). Interestingly, no other microorganisms of the community showed a phototrophic metabolism or presented the complete Calvin–Benson–Bassham (CBB) cycle. Only MAG 16 (*Methanosarcina mazei*) harboured a type III ribulose-1,5-bisphosphate carboxylase-oxygenase (RuBisCO) (Table 3; Table S6). Nevertheless, the type III RuBisCO participates in adenosine 5’-monophosphate (AMP) metabolism, a role that is distinct from that of classical RuBisCOs of the CBB cycle (Sato et al., 2007). MAG 1 (*Rhodopseudomonas faecalis)* has also a diazotrophic metabolism. It is able to fix N_2_ with a molybdenum-iron nitrogenase (Table 3; Table S7).

The enrichment of *R. faecalis* linked to the illumination of anaerobic digesters we report here could be further developed and used for the production of optimized biogas with a high CH_4_:CO_2_ ratio. In fact, a previous study reported an increment of methane production when the anaerobic digester operated under the influence of light, although a potential increment of *Rhosopseudomonas* species was not investigated by the authors (Tada et al., 2006). Although the biogas generated from anaerobic digestion processes is clean, carbon neutral, and environment-friendly, raw biogas often needs to be purified prior to its use. To date, several strategies have been designed to substantially reduce CO_2_ via chemical absorption (Akkarawatkhoosith et al., 2018). However, this process is expensive. Our findings support the possibility of biologically generating optimized biogas. Since *R. faecalis* has a photoautotrophic metabolism under anaerobic conditions, it can theoretically increase the CH_4_:CO_2_ ratio of the produced biogas through the fixation of CO_2_. The CH_4_:CO_2_ could be further increased by the action of iron-iron (Fe-only) nitrogenases, which have been recently reported to reduce CO_2_ simultaneously with nitrogen gas and protons to yield CH_4_, ammonia and hydrogen gas in a single enzymatic step (Zheng et al., 2018). Even though no iron-iron nitrogenases were detected in MAG 1 (*R. faecalis*) or in the whole metagenome, other *Rhodopseudomonas* strains harboring those enzymes could be useful to generate optimized biogas (Zheng et al., 2018).

Since the enrichment of *R. faecalis* in anaerobic digesters is expected to be dependent on the amount of light that can pass through the wall of the reactor, as well as on the ratio surface:volume of the reactor, the specific design and of illuminated anaerobic digesters is a critical and yet complex issue. Our results pave the way for future research aiming at optimising the development of *R. faecalis* light-dependent biofilms in order to optimize methane-rich biogas production in full-scale reactors.

## 4. Conclusions and further considerations

In the present work, we have carried out a complete study of a biofilm developing on the transparent wall of a lab-scale anaerobic digester operated under sunlight conditions. The microbial community harbored members involved in the four metabolic stages needed for the anaerobic digestion of organic matter, namely breakdown of polymers into monomers, acidification, acetogenesis and methanogenesis. *Methanosarcina* was the dominant methanogen in the anaerobic digester. The key difference with regard to conventional bioreactors that operate in darkness was a very significant enrichment of *R. faecalis*, a purple non-sulfur bacterium with a photoautotroph metabolism under anaerobic conditions. The ability of this bacterium to assimilate carbon dioxide through the CBB cycle, and its compatibility with the biogas process as well as with the rest of the microbiome opens up the striking possibility of producing optimized biogas from biomass through specifically designed, illuminated reactors.

## Supporting information

Captions of the Supplementary Tables

## Acknowledgements

We are thankful for the funding received from the German Ministry of Economic Affairs and Energy (grant no. 16KN070128, 16KN070126). Moreover, we also thank Justus Hardegen, Andreas Underberg and Anja Schmidt for technical assistance. Adriel Latorre is a recipient of a Doctorado Industrial fellowship from the Ministerio de Ciencia, Innovación y Universidades (reference DI-17-09613).

