## Supplementary material for "Shedding light on biogas: a transparent reactor triggers the development of a biofilm dominated by *Rhodopseudomonas faecalis* that holds potential for improved biogas production": Captions of the Supplementary Tables

**Table S1.** Taxonomic classification of bacterial 16S rRNA reads sequenced with the MinION.

**Table S2.** List of gene used with the UBCG v. 3.0 pipeline to generate the phylogenomic tree for each MAG. For each gene is indicated the number of nucleotides used.

**Table S3.** Classification of metagenomic coding sequences (CDS) into functional subsystems. For each subsystem the number of genes and roles are specified. It is distinguished between “active” and “likely” variant for each subsystem.

**Table S4.** Number of different carbohydrate-active enzymes (CAZyme) detected in each MAG.

**Table S5.** Presence key enzymes involved in fermetative and acetogenesis processes in each MAG. For each enzyme the number of orthologous genes is indicated.

**Table S6.** Enzymes involved in important metabolic processes detected in *Methanosarcina mazei* (MAG 16).

**Table S6.** Enzymes involved in important metabolic processes detected in *Methanosarcina mazei* (MAG 16). Patric ID were used to locate genes on the genome.

**Table S7.** Enzymes involved in important metabolic processes detected in *Rhodopseudomonas faecalis* (MAG 1). Patric ID were used to locate genes on the genome.
