## Supplementary material for "Shedding light on biogas: a transparent reactor triggers the development of a biofilm dominated by *Rhodopseudomonas faecalis* that holds potential for improved biogas production": Fig. S2

### Slide 1
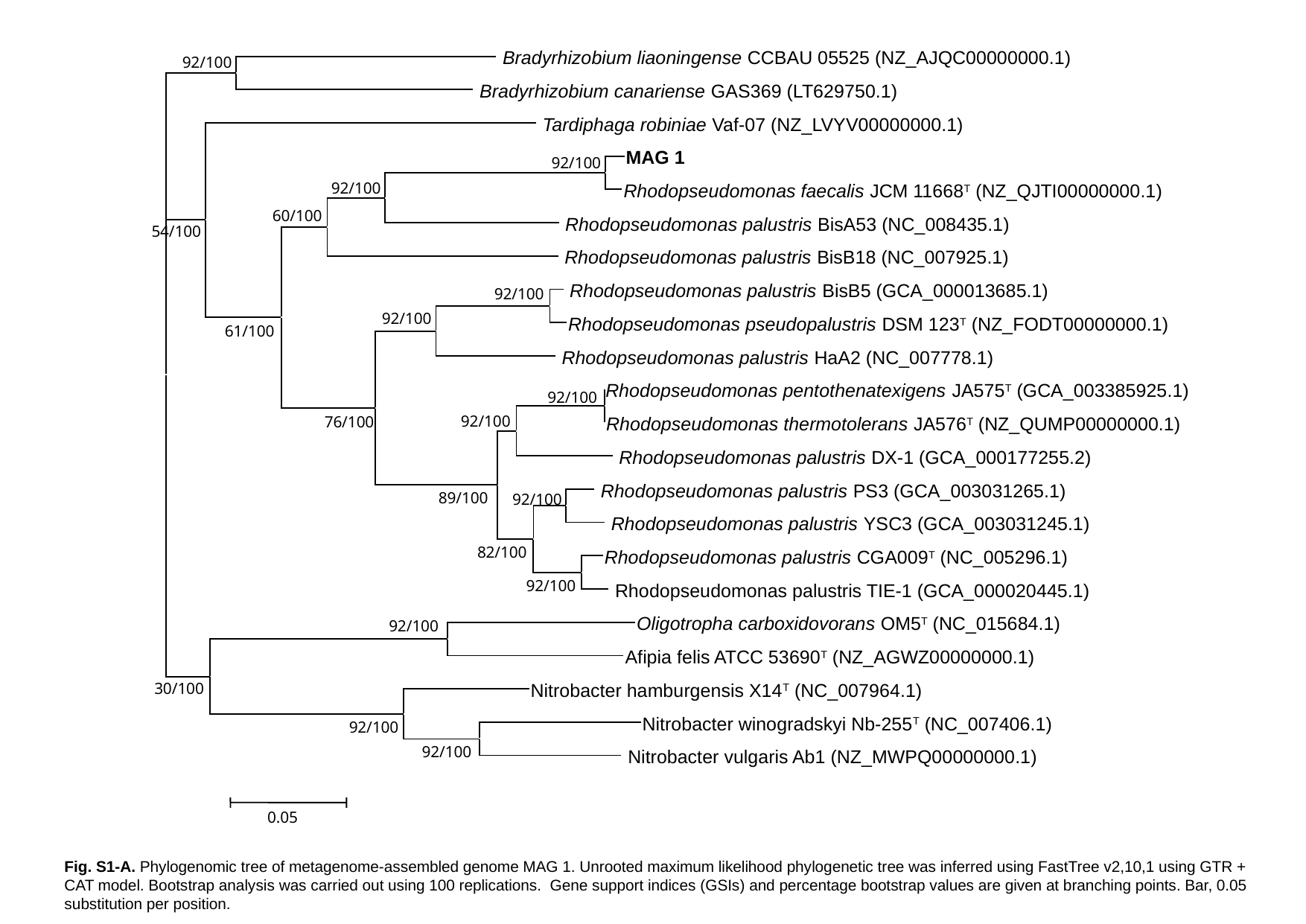

Bradyrhizobium liaoningense CCBAU 05525 (NZ_AJQC00000000.1)
92/100
 Bradyrhizobium canariense GAS369 (LT629750.1)
 Tardiphaga robiniae Vaf-07 (NZ_LVYV00000000.1)
MAG 1
92/100
92/100
Rhodopseudomonas faecalis JCM 11668^T^ (NZ_QJTI00000000.1)
60/100
 Rhodopseudomonas palustris BisA53 (NC_008435.1)
54/100
 Rhodopseudomonas palustris BisB18 (NC_007925.1)
 Rhodopseudomonas palustris BisB5 (GCA_000013685.1)
92/100
92/100
Rhodopseudomonas pseudopalustris DSM 123^T^ (NZ_FODT00000000.1)
61/100
 Rhodopseudomonas palustris HaA2 (NC_007778.1)
Rhodopseudomonas pentothenatexigens JA575^T^ (GCA_003385925.1)
92/100
92/100
76/100
Rhodopseudomonas thermotolerans JA576^T^ (NZ_QUMP00000000.1)
 Rhodopseudomonas palustris DX-1 (GCA_000177255.2)
 Rhodopseudomonas palustris PS3 (GCA_003031265.1)
89/100
92/100
 Rhodopseudomonas palustris YSC3 (GCA_003031245.1)
82/100
Rhodopseudomonas palustris CGA009^T^ (NC_005296.1)
92/100
 Rhodopseudomonas palustris TIE-1 (GCA_000020445.1)
Oligotropha carboxidovorans OM5^T^ (NC_015684.1)
92/100
Afipia felis ATCC 53690^T^ (NZ_AGWZ00000000.1)
30/100
Nitrobacter hamburgensis X14^T^ (NC_007964.1)
Nitrobacter winogradskyi Nb-255^T^ (NC_007406.1)
92/100
92/100
 Nitrobacter vulgaris Ab1 (NZ_MWPQ00000000.1)
0.05
Fig. S1-A. Phylogenomic tree of metagenome-assembled genome MAG 1. Unrooted maximum likelihood phylogenetic tree was inferred using FastTree v2,10,1 using GTR + CAT model. Bootstrap analysis was carried out using 100 replications. Gene support indices (GSIs) and percentage bootstrap values are given at branching points. Bar, 0.05 substitution per position.

### Slide 2
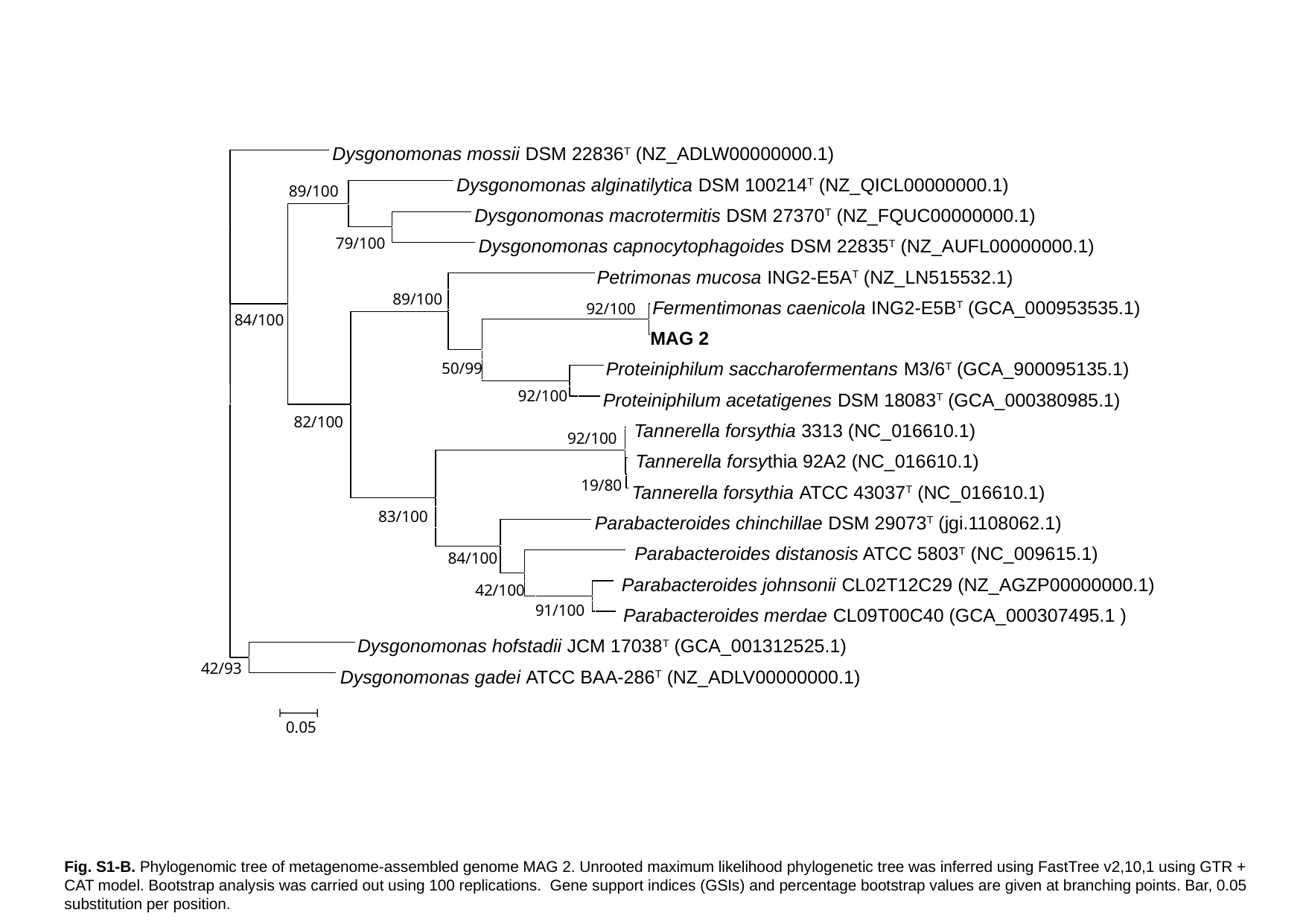

Dysgonomonas mossii DSM 22836^T^ (NZ_ADLW00000000.1)
Dysgonomonas alginatilytica DSM 100214^T^ (NZ_QICL00000000.1)
89/100
Dysgonomonas macrotermitis DSM 27370^T^ (NZ_FQUC00000000.1)
79/100
Dysgonomonas capnocytophagoides DSM 22835^T^ (NZ_AUFL00000000.1)
Petrimonas mucosa ING2-E5A^T^ (NZ_LN515532.1)
89/100
Fermentimonas caenicola ING2-E5B^T^ (GCA_000953535.1)
92/100
84/100
MAG 2
Proteiniphilum saccharofermentans M3/6^T^ (GCA_900095135.1)
50/99
92/100
Proteiniphilum acetatigenes DSM 18083^T^ (GCA_000380985.1)
82/100
 Tannerella forsythia 3313 (NC_016610.1)
92/100
 Tannerella forsythia 92A2 (NC_016610.1)
19/80
Tannerella forsythia ATCC 43037^T^ (NC_016610.1)
83/100
Parabacteroides chinchillae DSM 29073^T^ (jgi.1108062.1)
 Parabacteroides distanosis ATCC 5803^T^ (NC_009615.1)
84/100
 Parabacteroides johnsonii CL02T12C29 (NZ_AGZP00000000.1)
42/100
91/100
 Parabacteroides merdae CL09T00C40 (GCA_000307495.1 )
Dysgonomonas hofstadii JCM 17038^T^ (GCA_001312525.1)
42/93
Dysgonomonas gadei ATCC BAA-286^T^ (NZ_ADLV00000000.1)
0.05
Fig. S1-B. Phylogenomic tree of metagenome-assembled genome MAG 2. Unrooted maximum likelihood phylogenetic tree was inferred using FastTree v2,10,1 using GTR + CAT model. Bootstrap analysis was carried out using 100 replications. Gene support indices (GSIs) and percentage bootstrap values are given at branching points. Bar, 0.05 substitution per position.

### Slide 3
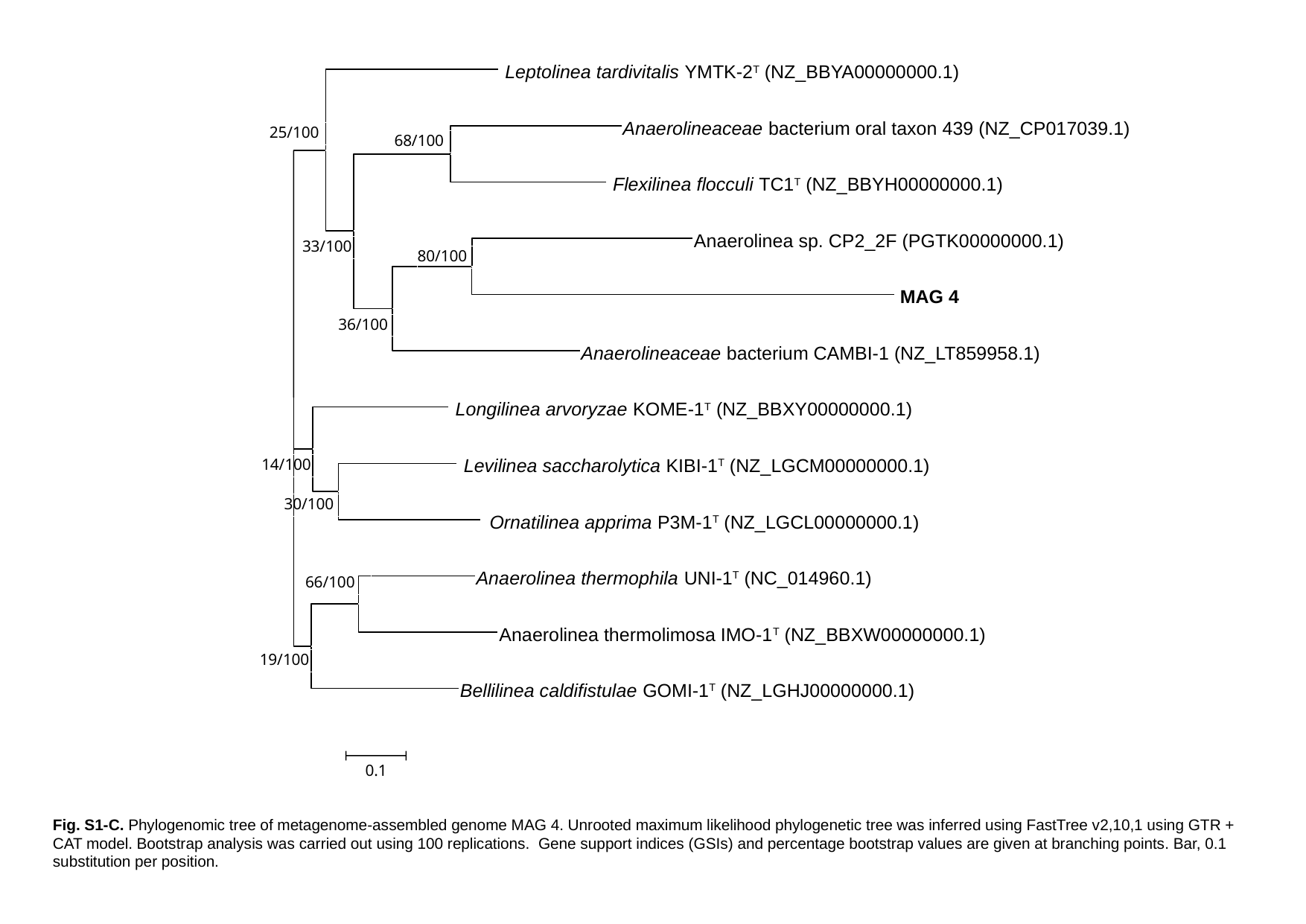

Leptolinea tardivitalis YMTK-2^T^ (NZ_BBYA00000000.1)
Anaerolineaceae bacterium oral taxon 439 (NZ_CP017039.1)
25/100
68/100
 Flexilinea flocculi TC1^T^ (NZ_BBYH00000000.1)
Anaerolinea sp. CP2_2F (PGTK00000000.1)
33/100
80/100
 MAG 4
36/100
Anaerolineaceae bacterium CAMBI-1 (NZ_LT859958.1)
 Longilinea arvoryzae KOME-1^T^ (NZ_BBXY00000000.1)
 Levilinea saccharolytica KIBI-1^T^ (NZ_LGCM00000000.1)
14/100
30/100
 Ornatilinea apprima P3M-1^T^ (NZ_LGCL00000000.1)
Anaerolinea thermophila UNI-1^T^ (NC_014960.1)
66/100
Anaerolinea thermolimosa IMO-1^T^ (NZ_BBXW00000000.1)
19/100
Bellilinea caldifistulae GOMI-1^T^ (NZ_LGHJ00000000.1)
0.1
Fig. S1-C. Phylogenomic tree of metagenome-assembled genome MAG 4. Unrooted maximum likelihood phylogenetic tree was inferred using FastTree v2,10,1 using GTR + CAT model. Bootstrap analysis was carried out using 100 replications. Gene support indices (GSIs) and percentage bootstrap values are given at branching points. Bar, 0.1 substitution per position.

### Slide 4
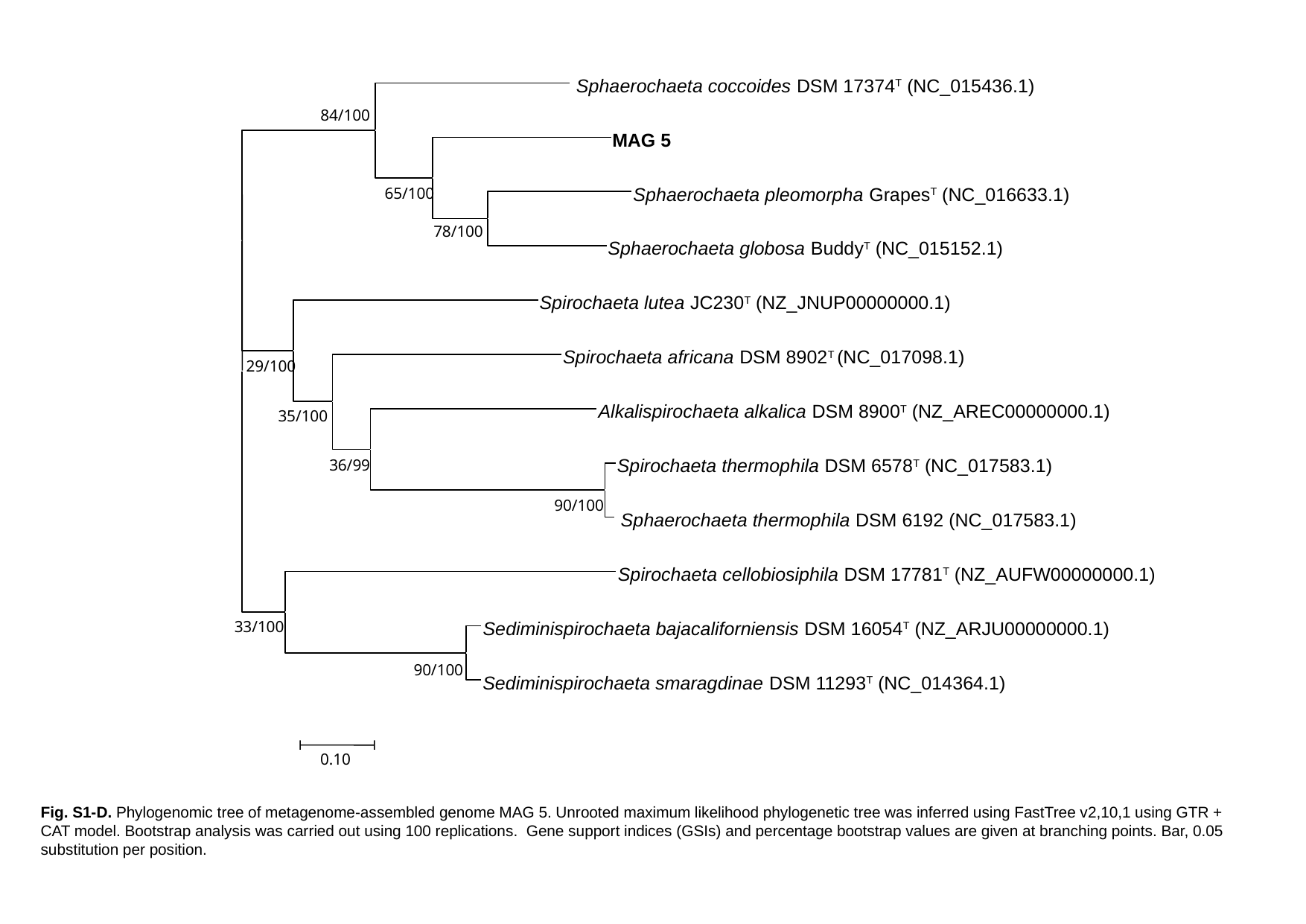

Sphaerochaeta coccoides DSM 17374^T^ (NC_015436.1)
84/100
MAG 5
Sphaerochaeta pleomorpha Grapes^T^ (NC_016633.1)
65/100
78/100
Sphaerochaeta globosa Buddy^T^ (NC_015152.1)
Spirochaeta lutea JC230^T^ (NZ_JNUP00000000.1)
Spirochaeta africana DSM 8902^T^ (NC_017098.1)
29/100
Alkalispirochaeta alkalica DSM 8900^T^ (NZ_AREC00000000.1)
35/100
Spirochaeta thermophila DSM 6578^T^ (NC_017583.1)
36/99
90/100
 Sphaerochaeta thermophila DSM 6192 (NC_017583.1)
Spirochaeta cellobiosiphila DSM 17781^T^ (NZ_AUFW00000000.1)
33/100
Sediminispirochaeta bajacaliforniensis DSM 16054^T^ (NZ_ARJU00000000.1)
90/100
Sediminispirochaeta smaragdinae DSM 11293^T^ (NC_014364.1)
0.10
Fig. S1-D. Phylogenomic tree of metagenome-assembled genome MAG 5. Unrooted maximum likelihood phylogenetic tree was inferred using FastTree v2,10,1 using GTR + CAT model. Bootstrap analysis was carried out using 100 replications. Gene support indices (GSIs) and percentage bootstrap values are given at branching points. Bar, 0.05 substitution per position.

### Slide 5
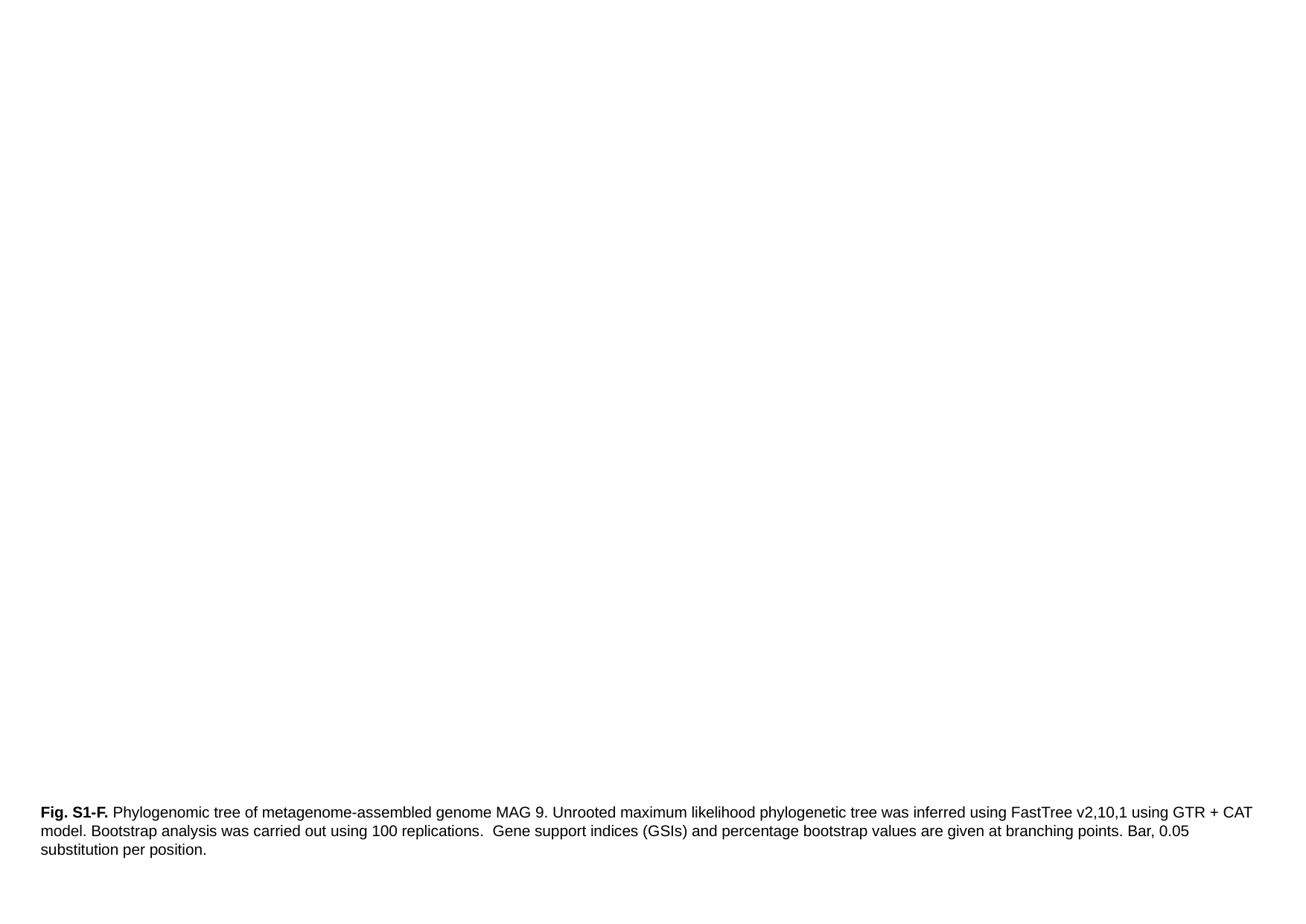

Fig. S1-F. Phylogenomic tree of metagenome-assembled genome MAG 9. Unrooted maximum likelihood phylogenetic tree was inferred using FastTree v2,10,1 using GTR + CAT model. Bootstrap analysis was carried out using 100 replications. Gene support indices (GSIs) and percentage bootstrap values are given at branching points. Bar, 0.05 substitution per position.

### Slide 6
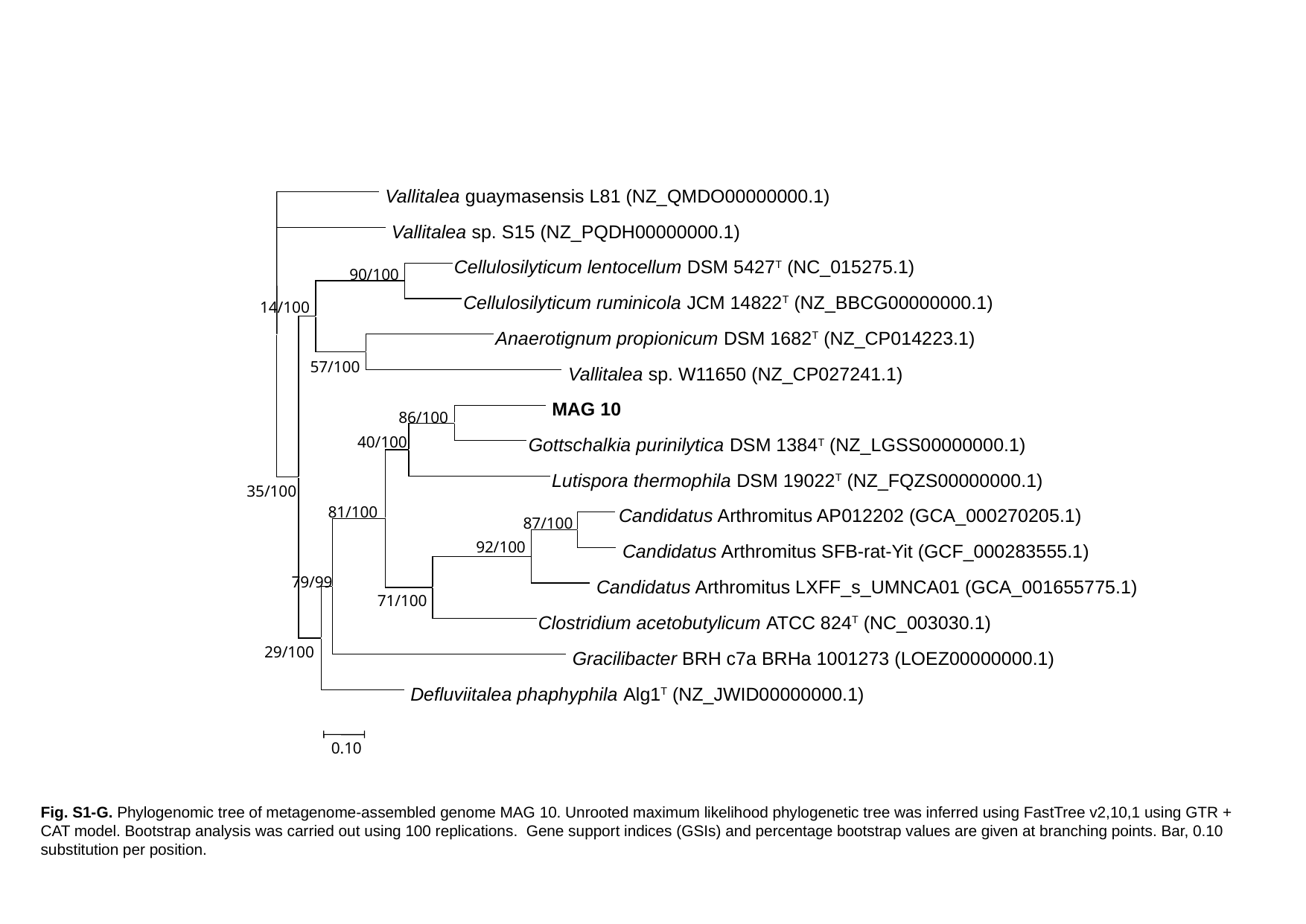

Vallitalea guaymasensis L81 (NZ_QMDO00000000.1)
 Vallitalea sp. S15 (NZ_PQDH00000000.1)
Cellulosilyticum lentocellum DSM 5427^T^ (NC_015275.1)
90/100
Cellulosilyticum ruminicola JCM 14822^T^ (NZ_BBCG00000000.1)
14/100
Anaerotignum propionicum DSM 1682^T^ (NZ_CP014223.1)
57/100
 Vallitalea sp. W11650 (NZ_CP027241.1)
 MAG 10
86/100
40/100
Gottschalkia purinilytica DSM 1384^T^ (NZ_LGSS00000000.1)
Lutispora thermophila DSM 19022^T^ (NZ_FQZS00000000.1)
35/100
81/100
Candidatus Arthromitus AP012202 (GCA_000270205.1)
87/100
92/100
 Candidatus Arthromitus SFB-rat-Yit (GCF_000283555.1)
79/99
 Candidatus Arthromitus LXFF_s_UMNCA01 (GCA_001655775.1)
71/100
Clostridium acetobutylicum ATCC 824^T^ (NC_003030.1)
29/100
 Gracilibacter BRH c7a BRHa 1001273 (LOEZ00000000.1)
 Defluviitalea phaphyphila Alg1^T^ (NZ_JWID00000000.1)
0.10
Fig. S1-G. Phylogenomic tree of metagenome-assembled genome MAG 10. Unrooted maximum likelihood phylogenetic tree was inferred using FastTree v2,10,1 using GTR + CAT model. Bootstrap analysis was carried out using 100 replications. Gene support indices (GSIs) and percentage bootstrap values are given at branching points. Bar, 0.10 substitution per position.

### Slide 7
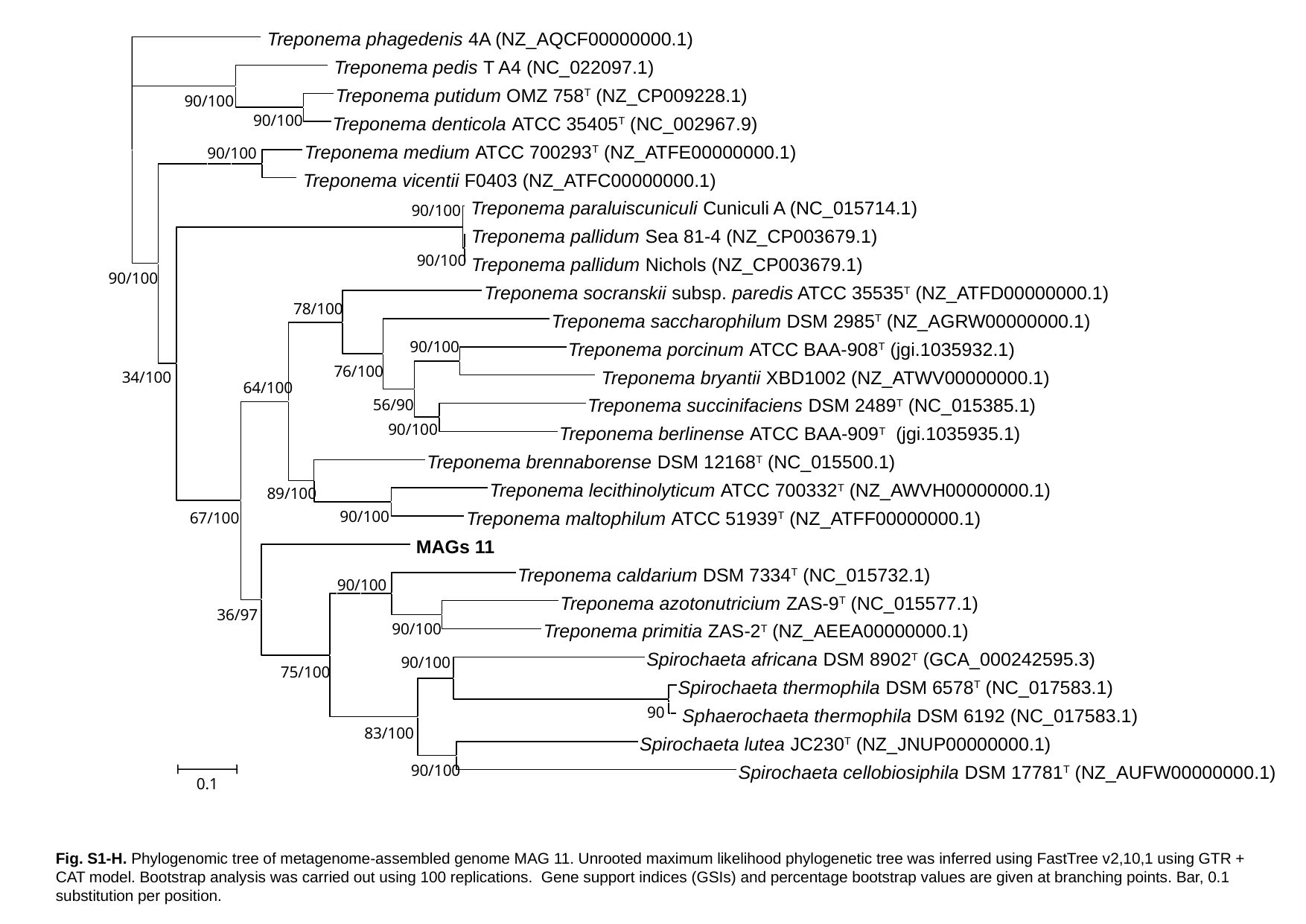

Treponema phagedenis 4A (NZ_AQCF00000000.1)
 Treponema pedis T A4 (NC_022097.1)
Treponema putidum OMZ 758^T^ (NZ_CP009228.1)
90/100
90/100
Treponema denticola ATCC 35405^T^ (NC_002967.9)
Treponema medium ATCC 700293^T^ (NZ_ATFE00000000.1)
90/100
 Treponema vicentii F0403 (NZ_ATFC00000000.1)
 Treponema paraluiscuniculi Cuniculi A (NC_015714.1)
90/100
 Treponema pallidum Sea 81-4 (NZ_CP003679.1)
90/100
 Treponema pallidum Nichols (NZ_CP003679.1)
90/100
Treponema socranskii subsp. paredis ATCC 35535^T^ (NZ_ATFD00000000.1)
78/100
Treponema saccharophilum DSM 2985^T^ (NZ_AGRW00000000.1)
90/100
Treponema porcinum ATCC BAA-908^T^ (jgi.1035932.1)
76/100
 Treponema bryantii XBD1002 (NZ_ATWV00000000.1)
34/100
64/100
Treponema succinifaciens DSM 2489^T^ (NC_015385.1)
56/90
90/100
Treponema berlinense ATCC BAA-909^T^ (jgi.1035935.1)
Treponema brennaborense DSM 12168^T^ (NC_015500.1)
Treponema lecithinolyticum ATCC 700332^T^ (NZ_AWVH00000000.1)
89/100
Treponema maltophilum ATCC 51939^T^ (NZ_ATFF00000000.1)
90/100
67/100
 MAGs 11
Treponema caldarium DSM 7334^T^ (NC_015732.1)
90/100
Treponema azotonutricium ZAS-9^T^ (NC_015577.1)
36/97
90/100
Treponema primitia ZAS-2^T^ (NZ_AEEA00000000.1)
Spirochaeta africana DSM 8902^T^ (GCA_000242595.3)
90/100
75/100
Spirochaeta thermophila DSM 6578^T^ (NC_017583.1)
90
 Sphaerochaeta thermophila DSM 6192 (NC_017583.1)
83/100
Spirochaeta lutea JC230^T^ (NZ_JNUP00000000.1)
Spirochaeta cellobiosiphila DSM 17781^T^ (NZ_AUFW00000000.1)
90/100
0.1
Fig. S1-H. Phylogenomic tree of metagenome-assembled genome MAG 11. Unrooted maximum likelihood phylogenetic tree was inferred using FastTree v2,10,1 using GTR + CAT model. Bootstrap analysis was carried out using 100 replications. Gene support indices (GSIs) and percentage bootstrap values are given at branching points. Bar, 0.1 substitution per position.

### Slide 8
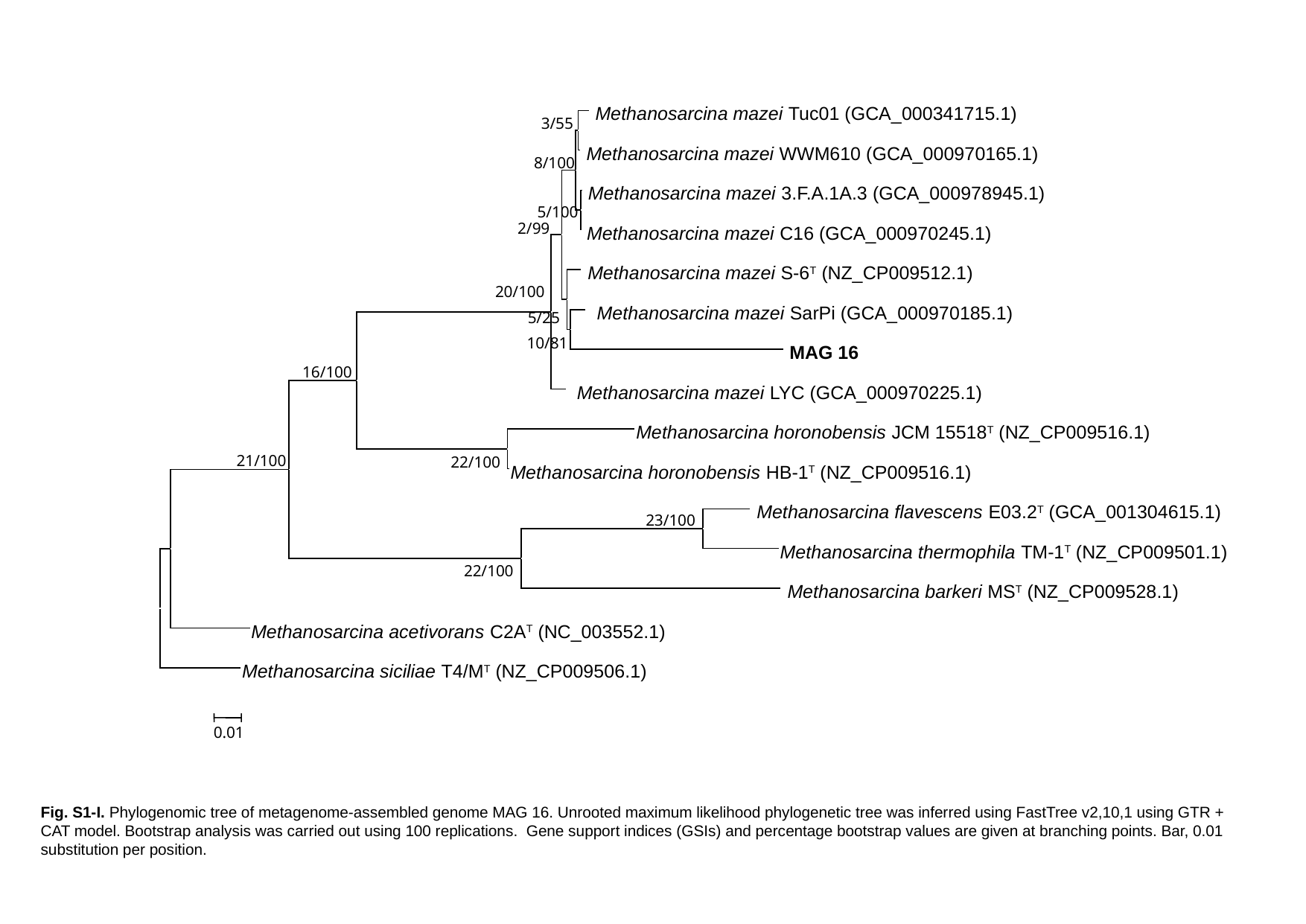

Methanosarcina mazei Tuc01 (GCA_000341715.1)
3/55
 Methanosarcina mazei WWM610 (GCA_000970165.1)
8/100
 Methanosarcina mazei 3.F.A.1A.3 (GCA_000978945.1)
5/100
2/99
 Methanosarcina mazei C16 (GCA_000970245.1)
 Methanosarcina mazei S-6^T^ (NZ_CP009512.1)
20/100
 Methanosarcina mazei SarPi (GCA_000970185.1)
5/25
10/81
 MAG 16
16/100
 Methanosarcina mazei LYC (GCA_000970225.1)
Methanosarcina horonobensis JCM 15518^T^ (NZ_CP009516.1)
21/100
22/100
Methanosarcina horonobensis HB-1^T^ (NZ_CP009516.1)
 Methanosarcina flavescens E03.2^T^ (GCA_001304615.1)
23/100
Methanosarcina thermophila TM-1^T^ (NZ_CP009501.1)
22/100
 Methanosarcina barkeri MS^T^ (NZ_CP009528.1)
Methanosarcina acetivorans C2A^T^ (NC_003552.1)
Methanosarcina siciliae T4/M^T^ (NZ_CP009506.1)
0.01
Fig. S1-I. Phylogenomic tree of metagenome-assembled genome MAG 16. Unrooted maximum likelihood phylogenetic tree was inferred using FastTree v2,10,1 using GTR + CAT model. Bootstrap analysis was carried out using 100 replications. Gene support indices (GSIs) and percentage bootstrap values are given at branching points. Bar, 0.01 substitution per position.
