## Supplementary material for "Shedding light on biogas: a transparent reactor triggers the development of a biofilm dominated by *Rhodopseudomonas faecalis* that holds potential for improved biogas production": Fig. S1

### Slide 1
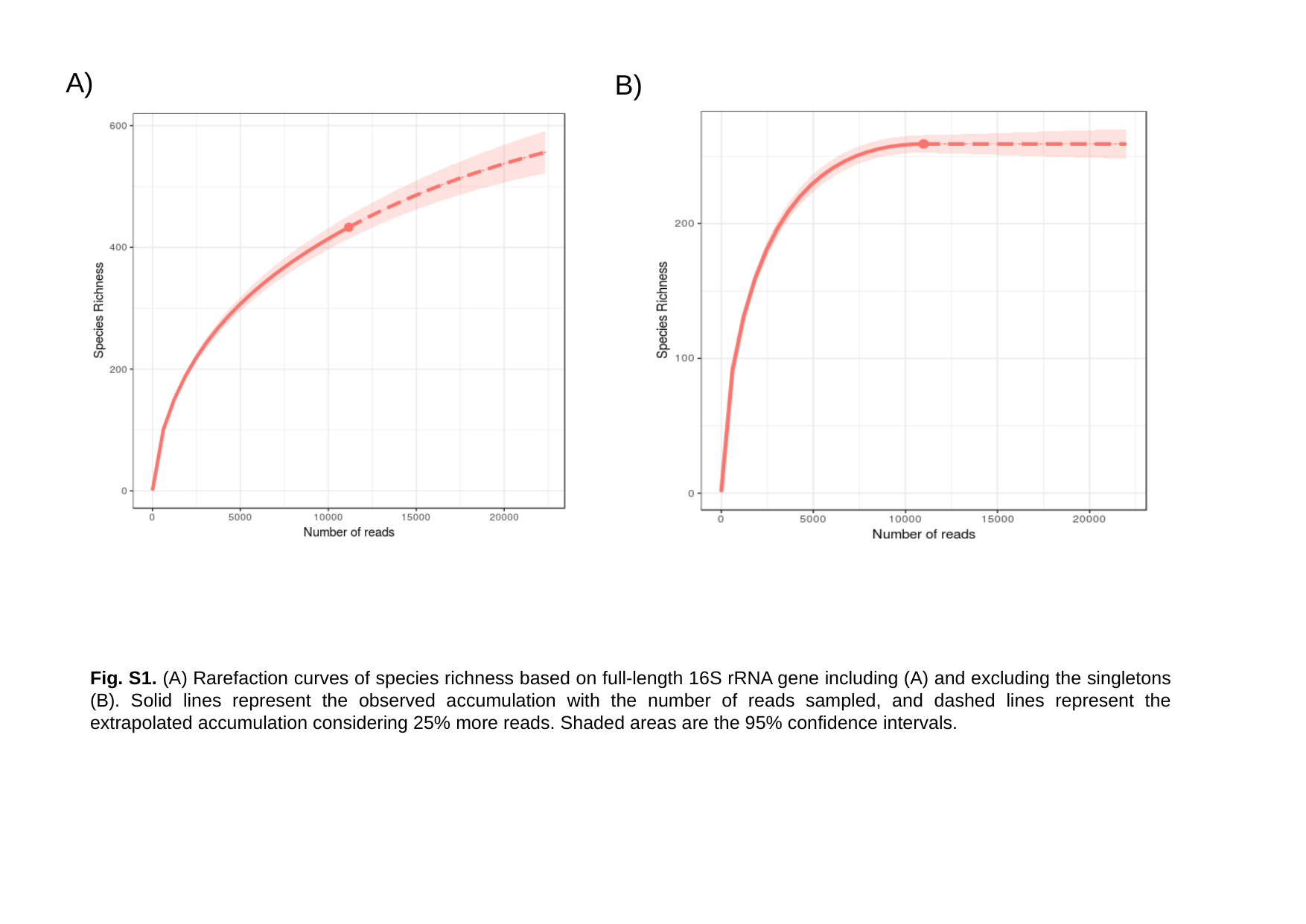

A)
B)
Fig. S1. (A) Rarefaction curves of species richness based on full-length 16S rRNA gene including (A) and excluding the singletons (B). Solid lines represent the observed accumulation with the number of reads sampled, and dashed lines represent the extrapolated accumulation considering 25% more reads. Shaded areas are the 95% confidence intervals.
